# Conserved induction of distinct antiviral signalling kinetics by primate interferon lambda 4 proteins

**DOI:** 10.1101/2021.09.15.460494

**Authors:** Cuncai Guo, Dorothee Reuss, Jonathon D. Coey, Swathi Sukumar, Benjamin Lang, John McLauchlan, Steeve Boulant, Megan L. Stanifer, Connor G. G. Bamford

**Affiliations:** Department of Infectious Diseases, Virology, University Hospital Heidelberg, Heidelberg, Germany; Wellcome-Wolfson Institute for Experimental Medicine, Queen’s University Belfast, Belfast, UK; Institute of Virology, University of Münster, Münster, Germany; Exzellenzcluster Science of Intelligence, Technische Universität Berlin, Berlin, Germany; Medical Research Council University of Glasgow Centre for Virus Research, University of Glasgow, Glasgow, UK; Research Group “Cellular Polarity and Viral Infection”, German Cancer Research Center (DKFZ), Heidelberg, Germany; Department of Molecular Genetics and Microbiology, College of Medicine, University of Florida, Gainesville, USA; Department of Infectious Diseases, Molecular Virology, University Hospital Heidelberg, Heidelberg, Germany

**Author notes:** co-corresponding authors **Contact details:** Megan Stanifer, Connor G G Bamford. These authors contributed equally (Cuncai Guo, Dorothee Reuss).

**Keywords:** interferon, lambda, signalling, antiviral, *IFNL4*, kinetics

## Abstract

Interferon lambdas (IFNλ) (also known as type III IFNs) are critical cytokines that combat infection predominantly at barrier tissues, such as the lung, liver and gastrointestinal tract. Humans have four IFNλs (1-4) where IFNλ1-3 show ∼80-95% homology and IFNλ4 is the most divergent displaying only ∼30% sequence identity. Variants in IFNλ4 in humans are associated with the outcome of infection, such as with hepatitis C virus. However, how IFNλ4 variants impact cytokine signalling in other tissues and how well this is conserved is largely unknown. In this study we address whether differences in antiviral signalling exist between IFNλ4 variants in human hepatocyte and intestinal cells, comparing them to IFNλ3. We demonstrate that compared to IFNλ3, wild-type human IFNλ4 induces a signalling response with distinct magnitudes and kinetics, which is modified by naturally-occurring variants P70S and K154E in both cell types. IFNλ4’s distinct antiviral response was more rapid yet transient compared to IFNλ1 and 3. Additionally, divergent antiviral kinetics were also observed using non-human primate IFNλs and cell lines. Furthermore, an IFNλ4-like receptor-interacting interface failed to alter IFNλ1’s kinetics. Together our data provide further evidence that major functional differences exist within the IFNλ gene family. These results highlight the possible tissue specialisation of IFNλs and encourage further investigation of the divergent, non-redundant activities of IFNλ4 and other IFNλs.

**Contribution to the Field:** Viral infections remain major causes of death and disease in humans and other animals. Interferons (IFNs) are a diverse group of host signalling proteins that can induce a potent antiviral state in cells and are intimately involved in the outcome of infection. Genetic variants within one IFN (interferon lambda 4, IFNλ4) are associated with the outcome of hepatitis C infection in humans. However, how IFNλ4 functions – and how natural variants affect its activity - remains poorly understood. Comparing how the antiviral activity changes over time following stimulation with different IFNλs, we identified that IFNλ4 induces a more rapid antiviral state compared to other IFNλs in liver and intestinal cells. Importantly, this response was conserved within human variants and between humans and non-human primates (chimpanzee and Rhesus macaque). Our results shed light on the unique functions of the divergent IFNλ4 protein.

## Introduction

Viral infections of mucosal surfaces like the lung, gut and liver (such as influenza, rotavirus and hepatitis C virus (HCV)) remain major drivers of global morbidity and mortality in the human population (Abbafati et al. 2020). The host innate immune response is a critical determinant of the outcome of infection and as such, its stimulation can influence clinical outcomes (Heim, 2013). Following sensing of viral infection, several antiviral and immunoregulatory factors like cytokines are induced that act to limit viral replication and promote clearance and long-term immunity (Bowie and Unterholzner 2008). Interferons (IFNs) are one important group of such cytokines with potent antiviral activity (Isaacs and Lindenmann 1957). There exist three recognised families of IFNs: the type I IFNs (alpha 1- 13, beta, epsilon, kappa and omega in humans), type II IFNs (gamma) and type III IFNs (lambdas [λ] 1-4) (Hoffmann, Schneider and Rice 2015). Types I and III IFNs are rapidly induced and secreted following sensing of infection in most nucleated cells. These secreted IFNs then act in turn on the infected cell as well as on neighbouring uninfected cells to induce the production of hundreds of interferon stimulated genes (ISGs) via activation of the JAK-STAT pathway. Although they share similar downstream signalling pathways and lead to the activation of similar ISGs, type I and III IFNs utilise distinct cell surface receptor complexes (Kotenko et al. 2003). Type I IFNs use the ubiquitously expressed IFNAR1 and IFNAR2 heterodimeric complex while type III IFNs use the IFNλR1 and IL10R2 heterodimeric complex. Although also found on some immune cell types (Santer et al. 2020), IFNλR1 is predominantly expressed on epithelial cells at so-called barrier tissues (Sommereyns et al. 2008b), including the respiratory and gastrointestinal tracts, as well as hepatocytes in the liver of humans (Sheppard et al. 2003), which provides type III IFNs distinct traits specialised in the protection of mucosal surfaces compared to type I IFNs (Marcello et al. 2006; Pervolaraki et al. 2018; Forero et al. 2019).

Although they share a receptor complex there is emerging evidence that not all type III IFNs have redundant features(Prokunina-Olsson et al. 2013). The human IFNλs: IFNλ1, IFNλ2, IFNλ3 all share >80% homology yet compared to IFNλ4 exhibit only ∼30% homology (Kotenko et al. 2003; Sheppard et al. 2003; Prokunina-Olsson et al. 2013). While all type III IFNs are more recently discovered in comparison to type I IFNs (Isaacs and Lindenmann 1957; Sheppard et al. 2003; Kotenko et al. 2003), IFNλ4 was the latest addition to the family being only identified in 2013 (Prokunina-Olsson et al. 2013). The outcome of HCV infection is associated with genetic variation in the human *IFNL* locus (e.g. ‘*IL28B’* SNPs), likely mediated by variants within *IFNL4* (Terczyńska-Dyla et al. 2014). IFNλ4 like other IFNλs has potent antiviral activity (Hamming et al. 2013). These same genetic variants are also associated with extra-hepatic infections, such as enteroviral infection in the respiratory tract (Rugwizangoga et al. 2019). There are two common loss-of-function single nucleotide polymorphisms (SNPs) in human *IFNL4*, encoding a frameshift (rs12979860), and a non-synonymous variant P70S (rs117648444, which encodes a proline to serine mutation at position 70), respectively (Prokunina-Olsson et al. 2013; Terczyńska-Dyla et al. 2014). While the frameshift ablates IFNλ4 production, P70S reduces the potency of ISG induction by IFNλ4 (Terczyńska-Dyla et al. 2014; Bamford et al. 2018; Hong et al. 2016). Interestingly, it is these hypo- or inactive alleles that are associated with protection from chronic HCV infection in humans(Prokunina-Olsson et al. 2013; Terczyńska-Dyla et al. 2014).

Further investigation into the functional diversity of IFNλ4 identified two rare variants that affect IFNλ4 activity, including an additional hypoactive variant L79F (leucine to phenylalanine at position 79), and K154E (lysine to glutamic acid at position 154), which dramatically enhances IFNλ4 antiviral activity by increasing its secretion and potency (Bamford et al. 2018). Intriguingly, although K154 is nearly ubiquitous in the human population, E154 is the ancestral amino acid at this position in non-human primates and other mammals. E154 was found in a small number of extant humans. Accordingly, chimpanzee and Rhesus macaque IFNλ4 have enhanced antiviral activity relative to wild-type human IFNλ4, which can be reversed by an E154K mutation. Together, the evolutionary data suggest a step-wise attenuation of IFNλ4 activity (E154K > P70S > TT frameshift) unique to modern humans(Prokunina-Olsson et al. 2013), which is consistent with the non-redundancy of IFNλ4 compared to other IFNλs. However, which precise unique biological feature(s) of IFNλ4 that are non-redundant (and thus have been acted upon by evolution) are poorly understood and only beginning to be unravelled (Terczyńska-Dyla et al. 2014; Zhou et al. 2020; Obajemu et al. 2017).

Following on from our previous work (Bamford et al. 2018; Pervolaraki et al. 2018) we wished to determine how the antiviral activity of IFNλ4 and its variants and homologues changed in a time-dependent manner, compared to other IFNλs. To test this hypothesis we characterised the kinetics of signalling and antiviral activity of a panel of IFNλ4 variants in human hepatocyte and human intestinal epithelial cells compared to IFNλ3. Together, our work demonstrates the unique kinetics of IFNλ4 activity compared to other IFNλs, which is conserved within and between species. Further work on the intrinsic differences between IFNλ4 and other IFNs is warranted.

## Results

### IFNλ variants display unique STAT1 phosphorylation kinetics

Binding of IFNλs to their receptor complex leads to activation of downstream signalling cascades that ultimately lead to the establishment of an antiviral state (Kotenko et al. 2003). The Janus kinase/signal transducer and activator of transcription (JAK/STAT) pathway is one of the most critical and well characterized pathways activated following IFNλ binding. An emerging view is that the kinetics of such a downstream response is a crucial determinant of the antiviral potential of IFNλs (Pervolaraki et al. 2018; Obajemu et al. 2017). To probe the temporal basis of IFNλ signalling in greater detail, we first measured phosphorylation of STAT1 over time at Y701 (**Fig 1**). Human hepatocyte HepaRG monolayers were incubated with conditioned media estimated to contain equivalent amounts of IFNλs (IFNλ3, IFNλ4 WT, P70S, L79F and K154E) for 15, 30, 60, 120, 360 mins and 24h. Following stimulation, protein lysates were harvested and STAT1 phosphorylation was assayed by immunoblot analysis (**Fig 1A**). Conditioned media generated following transfection of an EGFP-expressing plasmid served as a negative control. Results showed that IFNλ3 and a number of IFNλ4 variants induced detectable levels of pSTAT1 (**Fig 1A**, quantified in **sFig1A**). L79F gave extremely low levels of pSTAT1 (data not shown) which likely correlates with its very limited activity as described previously (Bamford et al. 2018). Interestingly, IFNλ3 and IFNλ4 variants induced distinct kinetics of pSTAT1 activation (**Fig 1A and sFig 1A**). While all IFNλs peaked around similar times (30min to 1 hour), IFNλ4 WT and P70S showed clear transient activation while IFNλ3 and K154E displayed persistent activation of pSTAT1. Importantly, levels of pSTAT1 correlated with previously measured antiviral potential for three IFNλ4 variants K154E>WT>P70S (Bamford et al. 2018). As IFNλs can also signal in other tissues apart from the human liver (Sommereyns et al. 2008; Pervolaraki et al. 2017), and there is an emerging role for IFNλ4 in extra-hepatic environments, we assayed whether human colon carcinoma cells (T84) were capable of inducing pSTAT1 in response to IFNλ4 and its variants. Intestinal T84 cells were treated with incubated with conditioned media containing equivalent amounts of IFNλs (IFNλ3, IFNλ4 WT, P70S, and K154E) and their induction of pSTAT1 was assayed over time by immunoblot analysis (**Fig 1B and sFig 1B)**. We observed similar trends as to HepaRG although differences in amplitude of pSTAT1 induction were noted, especially for IFNλ4 K154E in T84 cells. Together these results show that both hepatic and intestinal cell lines can respond to both IFNλ3 and IFNλ4 and display variant specific inductions of the JAK/STAT pathway.

**Figure 1.**
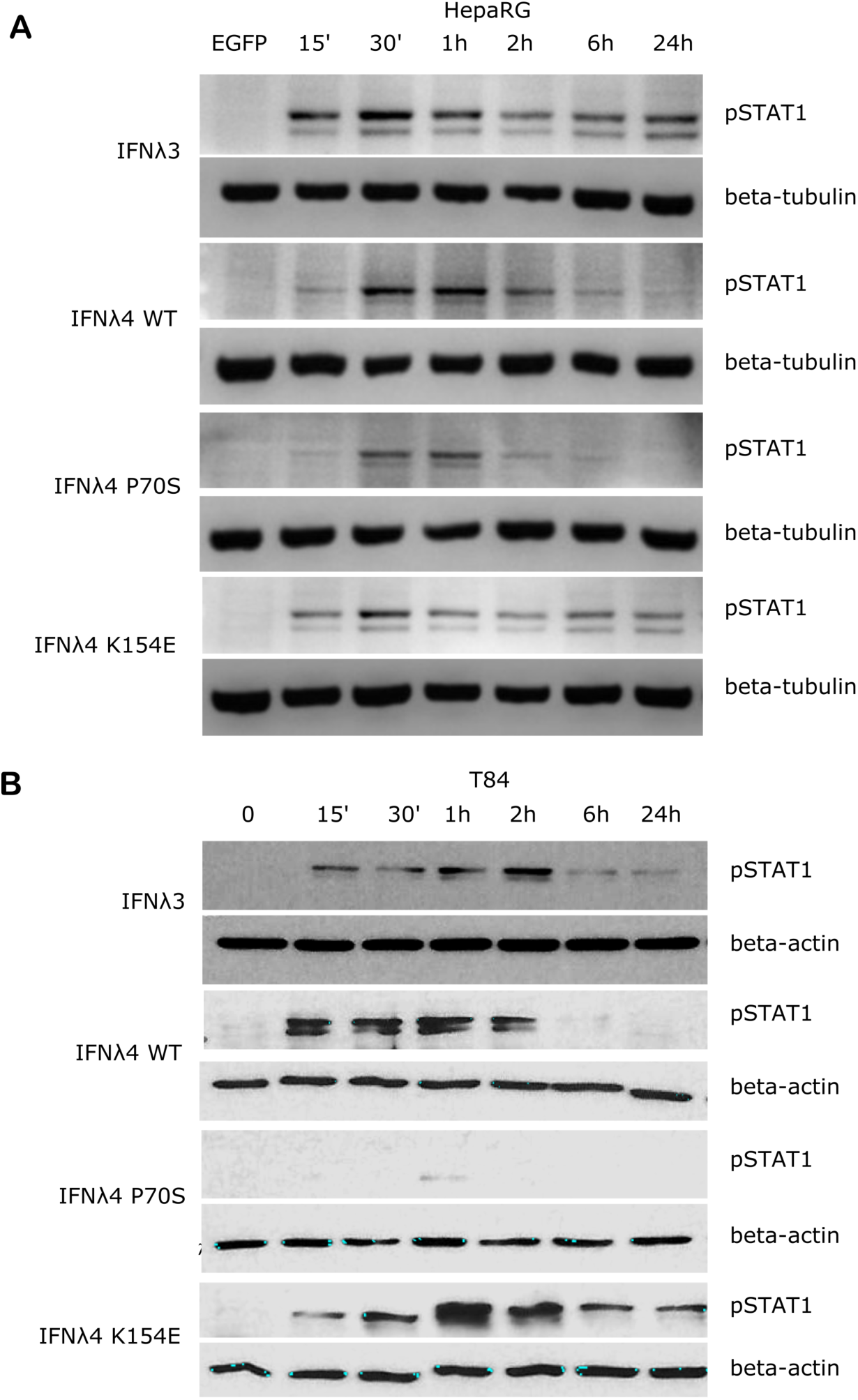
IFNλs each have a distinct kinetic of STAT1 phosphorylation. HepaRG (**A**) and T84 (**B**) cells were incubated with IFNλs (IFNλ3-HiBiT, IFNλ4-HiBiT: WT, P70S, and K154E for the indicated times and the levels of pSTAT1 was assayed by immunoblot. Beta-tublin (HepaRG) or beta-actin (T84) served as loading controls. EGFP (**A**) or timepoint 0 (**B**) serves as a conditioned media control. Representative images of 2-3 replicates are shown.

### IFNλ variants display different levels of ISG induction

Phosphorylation of STAT1 following receptor complex engagement by IFNλs results in STAT1/2 dimer formation and translocation to the nucleus to induce ISG transcription, which ultimately leads to the production of antiviral proteins and the establishment of an antiviral state (Kotenko et al. 2003). Our previous work showed that IFNλ variants induced different levels of ISG expression when measured at 24h(Bamford et al. 2018). To ascertain whether this ISG expression varied at earlier times after incubation in concert with the kinetics of pSTAT1 activation, we measured the relative induction of a panel of core ISGs (*IFIT1, MX1, ISG15*, and *RSAD2/VIPERIN*) compared to EGFP-treated conditioned media in HepaRG cells (**Fig 2A-D**) and T84 cells (**Fig 2E-H**). Compared to EGFP conditioned media stimulated cells, all IFNλs induced measurable increases in ISG mRNA in HepaRG cells but with discernible differences in magnitude. T84 cells also showed ISG induction for four of the supernatants tested (**Fig 2E-H**). Additionally, looking at relative fold change, T84 cells gave a lower induction of all ISGs as compared to HepaRG cells (**Fig 2**). The magnitudes of ISG induction for both cell lines mirrored the pSTAT1 induction that was observed in Fig. 1 (IFNλ3/K154E>WT>P70S>L79F). IFNλ4 K154E induced a similar pattern of ISG induction as IFNλ3 in both cell lines. Interestingly the kinetics of ISG induction was distinct to each cell line. In HepaRG cells, all IFNλs induced an early peak induction of ISGs, which subsequently declined over time. Moreover, IFNλ4 K154E demonstrated a slightly faster induction and peaked by 2h while all other IFNλs tested peaked at 6h. By contrast, IFNλ3 and the IFNλ4 K154E showed no or little decline in ISG induction after induction at either 2h or 6h in T84 cells (**Fig 2E-H**). Additionally, T84 cells yielded low to almost undetectable induction of ISGs following IFNλ4 WT and P70S treatment. Together these results show that K154E provides similar stimulatory activity to IFNλ3 and that this is far greater than for either IFNλ4 WT or P70S, which are the most common IFNλ4 variants in the human population.

**Figure 2.**
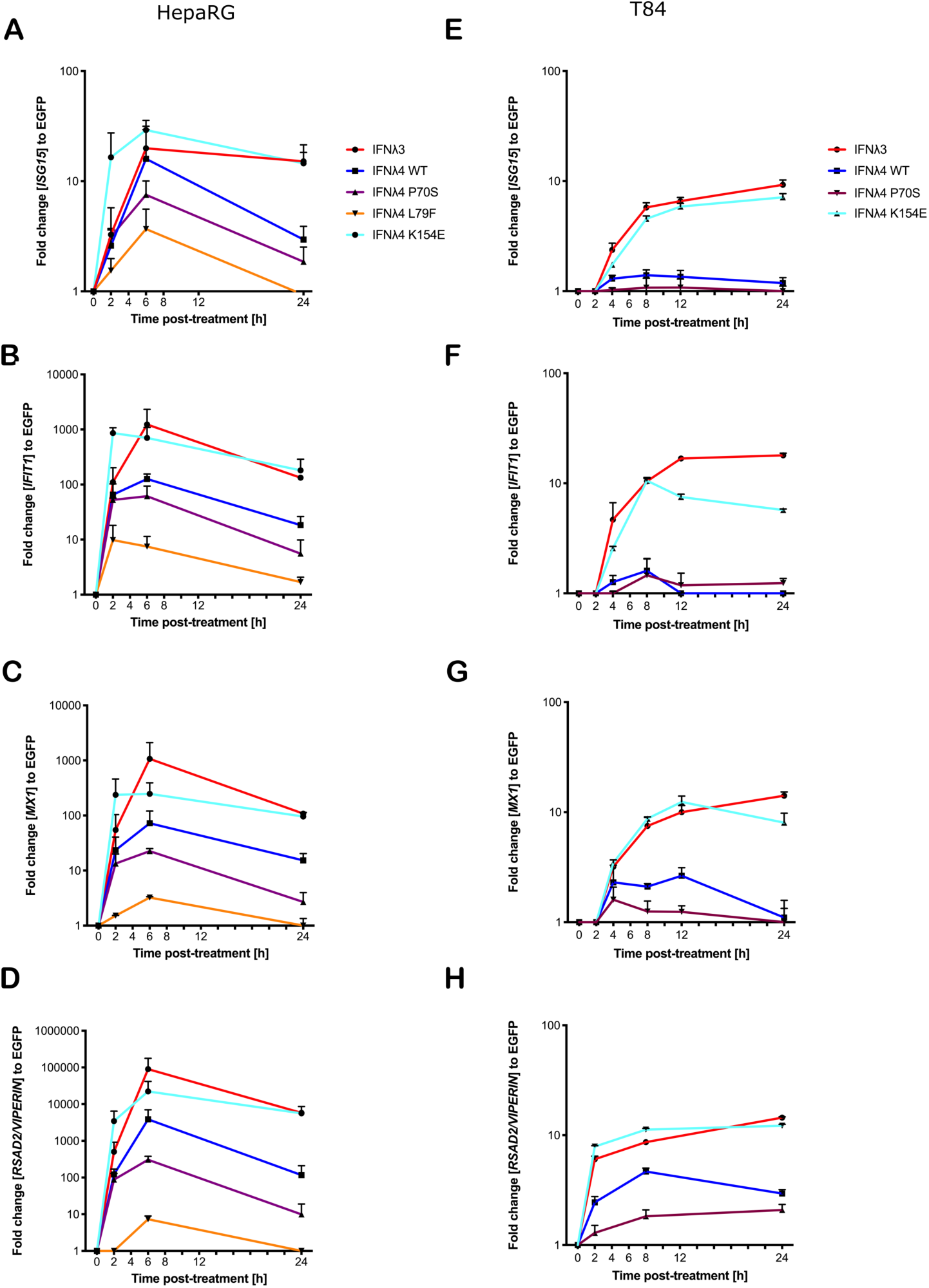
IFNλ variants induce unique magnitudes of ISG mRNA. HepaRG (**A-D**) and T84 (**E-H**) cells were incubated with IFNλs (IFNλ3-HiBiT [red], IFNλ4-HiBiT: WT [blue], P70S [purple], L79F [yellow] and K154E [cyan]) for indicated times. At the respective time, total RNA was isolated and qRT-PCR was performed for ISGs: *IFIT1* (**A and E**), *ISG15* (**B and F**), *MX1* (**C and G**) and *RSAD2/VIPERIN* (**D and H**). EGFP-treated cells were used as a mock control and all values were normalized against this value at each time. *GAPDH* (HepaRG) or *HPRT1* (T84 cells) were used as housekeeping genes. L79F did not induce any detectable ISG induction in T84 cells. Error bars represent the mean ± SEM from 2-3 biological replicates.

### IFNλ variants have distinct antiviral activity in intestinal cells

Induction of an antiviral state is the major downstream consequence of IFN signalling. To determine how STAT1 phosphorylation and ISG expression correlate with antiviral activity, we infected the hepatic and intestinal cell models with two different viruses, EMCV and VSV. Both EMCV and VSV are highly cytopathic, replicate very fast, and are sensitive to IFN which makes them suitable for assessing the kinetics of antiviral activity. EMCV infectivity and replication were assayed by determining the cytopathic effects of the virus while a VSV encoding luciferase (VSV-luc) was deployed and its infectivity was measured by luciferase assay. HepaRG and T84 cells were treated with increasing concentrations of EGFP or IFNλ3 or IFNλ4-containing supernatants at 24 hours prior to virus infection. Following IFNλ pre-treatment, cells were infected with EMCV or VSV (MOI of 0.3 and 1, respectively) in the continuous presence of IFNλs, and infection was assayed at 24h post-infection for EMCV (**Fig 3A**) and 8h post-infection for VSV (**Fig 3B and C**). Different assay times for VSV versus EMCV were due to differences in replication kinetics and cytopathic effects of either virus. Consistent with our previous work(Bamford et al. 2018; K et al. 2017), results show that VSV infection was inhibited by all IFNs in both cell lines (**Fig 3B and C**). IFNλ3 was the most potent IFN, as it reduced VSV infection with 10% of the maximum concentration in both HepRG and T84 cells. IFNλ4 WT and K154E showed similar antiviral activity however a much higher concentration of these two IFNs was required to reach a similar potency as IFNλ3. Consistent with previous low pSTAT1 and ISG inductions, P70S was only able to slightly reduce virus infection even at the highest concentrations in both cell lines. T84 cells were poorly infected with EMCV and highly resistant to the cytopathic effects of the virus and therefore, antiviral activity was not assayed in this cell line but similar patterns of antiviral activity were seen for HepaRG cells infected with EMCV (**Fig 3A**).

**Fig 3.**
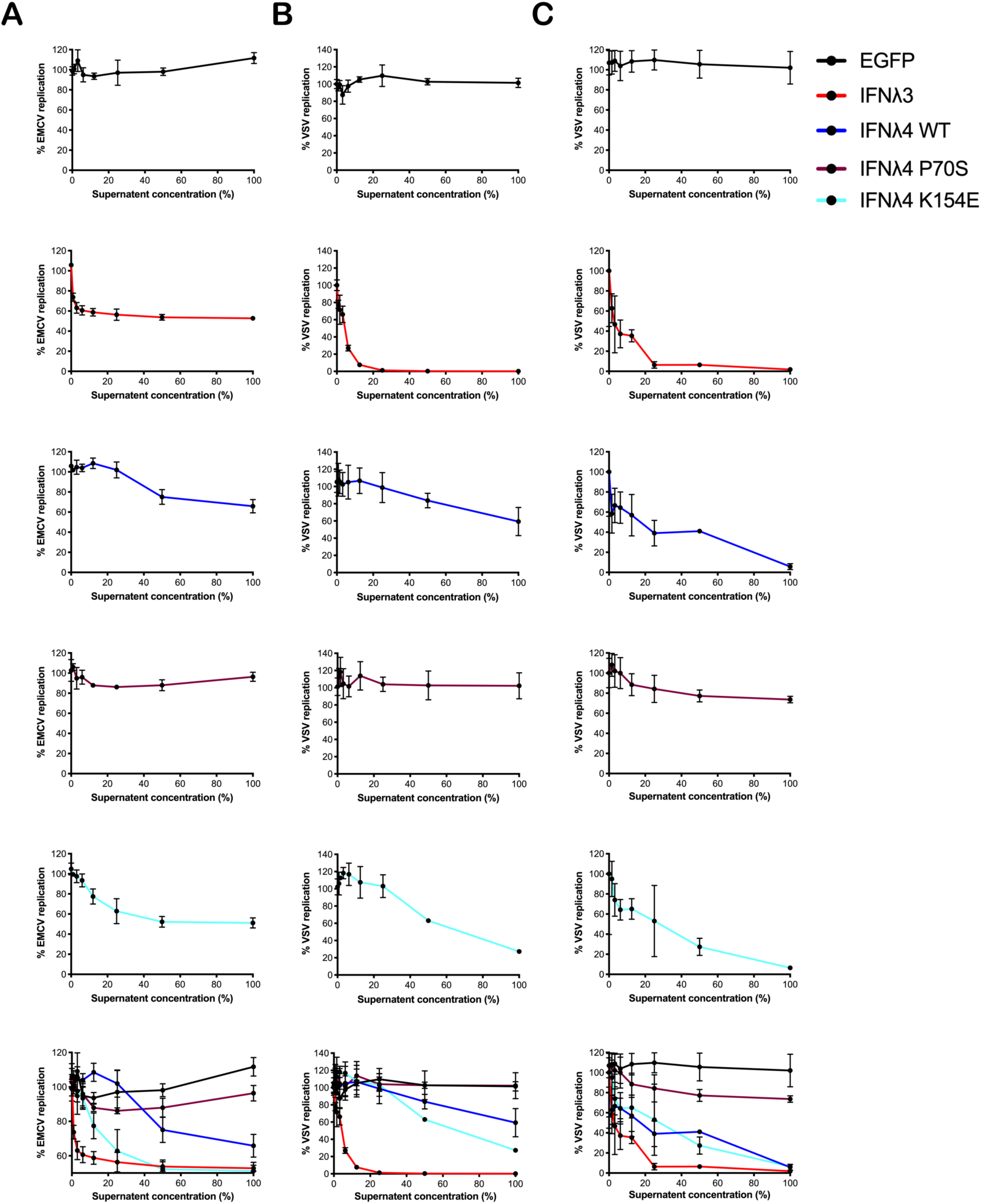
Antiviral activity against EMCV or VSV of IFNλs on HepaRG and T84 cells. HepaRG (**A and B**) or T84 (**C**) cells were stimulated with different concentrations of supernatant containing the panel of IFNλs (IFNλ3-HiBiT [red], IFNλ4-HiBiT: WT [blue], P70S [purple], and K154E [cyan]) before being challenged with EMCV (**A**) or VSV (**B and C**) and antiviral activity calculated, shown here as percentage of viral replication at each dilution compared to mock treated controls. Error bars represent the mean ± SEM from 2-3 biological replicates.

### IFNλ variants have distinct kinetics of antiviral activity

Having established antiviral assays in both liver- and intestinal-derived cell lines, we wished determine whether IFNλ activity was time-dependent, and whether the continuous presence of IFNλs was required to maintain their antiviral activity. Therefore, we performed infections and antiviral assays over time, both in the continuous presence of IFNλs but also in cells that had been pre-treated with IFNλs for varying lengths of time yet the cytokines were then removed, monolayers washed and fresh media provided (‘non-washed’ and ‘washed’ respectively, **Fig 4A**) prior to infection. Initially, we conducted experiments in T84 cells that were infected with VSV following IFNλ pre-treatment (**Fig 4B-E**). In agreement with the data presented in Fig 3, all IFNλs demonstrated antiviral activity with IFNλ3 and IFNλ4 P70S showing the greatest and least potency respectively. The peak of IFNλ3 activity was delayed relative to all IFNλ4s. Moreover, we found that shorter incubation times with IFNλ3 followed by its removal before infection reduced its antiviral activity to a greater extent compared to the three IFNλ4 variants used in the experiment (compare early time points in **Fig 4B** with **Figs 4C-E**). In HepaRG cells infected with EMCV, we observed a similar pattern, i.e removal of IFNλ3 after relatively short incubation (2h) with cells gave a greater reduction in antiviral activity compared to the same timepoint for the IFNλ4 variants (**Fig 4F-I**). In addition, we observed that all IFNλs generally gave less antiviral activity after removal at the time of infection compared to activities in the continuous presence of the proteins. From these experiments, we suggest that IFNλ4 proteins may be more tightly bound to the heteromeric cell receptor as compared to IFNλ3. Alternatively, signalling with IFNλ4 is maintained for a longer period as compared to IFNλ3. To further assess the contribution of IFN-cell contact time compared to signalling time, we repeated the wash experiments in HepaRG cells with EMCV but, on this occasion, cells were incubated with the IFNλs at 24h prior to infection but then removed by washing at differing times before virus addition (**sFig 2**). The results show a greater decline in antiviral activity (∼15-fold) from 24h incubation to 6 h and 2h incubation for IFNλ3 compared to IFNλ4 WT and IFNλ4 K154E which showed reductions in activity by ∼0.75-4 fold.

**Figure 4.**
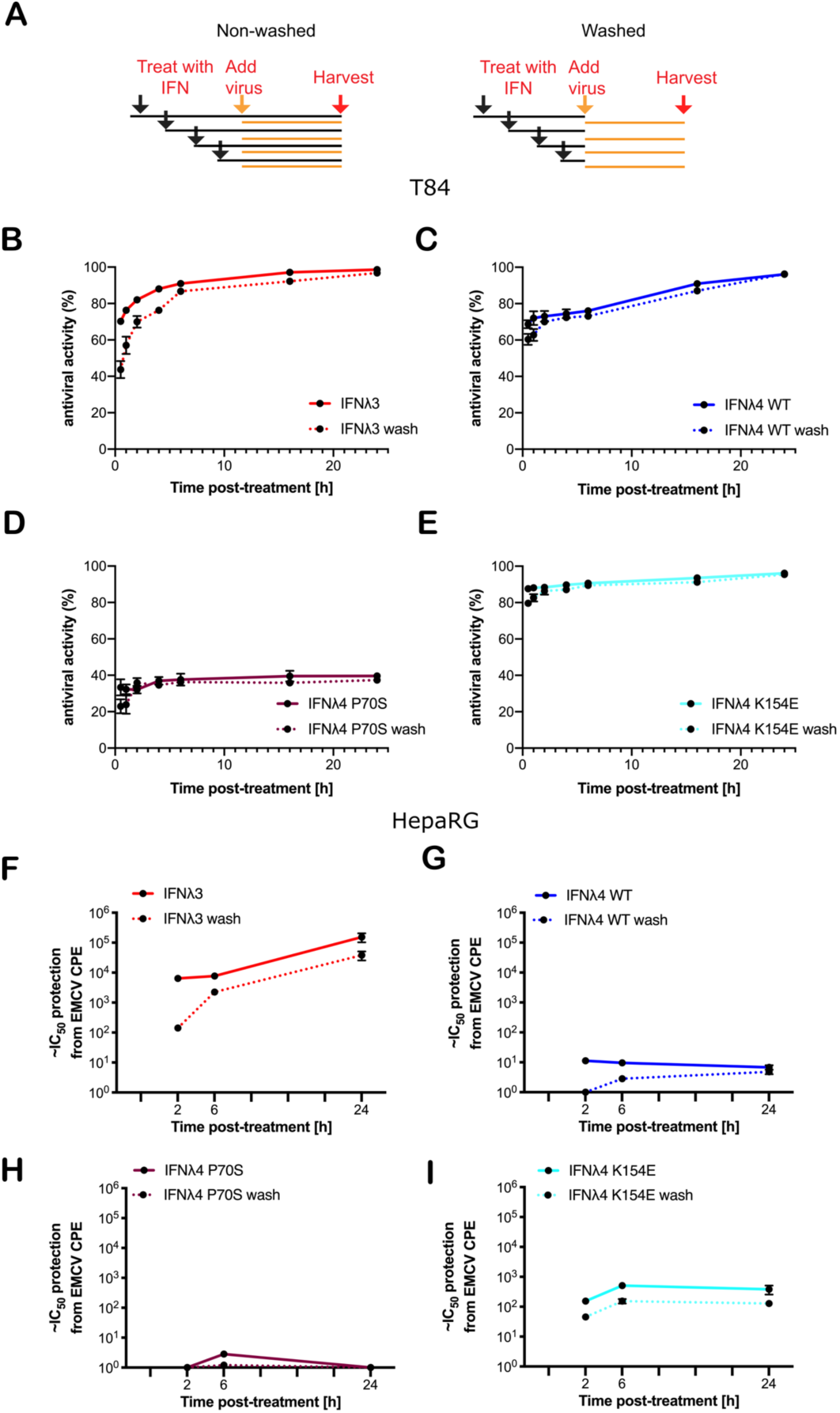
Antiviral activity does not require continued presence of IFNλs. **(A)** Schematic description of the experiment to show how IFNλ was added and maintained or removed by washing. T84 (**B-E**) and HepaRG (**F-I**) were stimulated with IFNλs: IFNλ3-HiBiT (**B and F**), IFNλ4-HiBiT WT (**C and G**), P70S (**D and H**), and K154E (**E and I**) at indicated time prior to infection with EMCV (HepaRG) or VSV (T84). VSV-luc (**B-E**) was assayed 8 hours post-infection by quantifying the luminescence (T84). EMCV infection (**F-I**) was assayed by analysis of its cytopathic effect 24 hours post-infection of a series of two-fold serial dilutions of supernatant. For washing experiments (dashed lines), IFNλs were removed and rinsed with PBS before being replaced with media containing virus. Error bars represent the mean ± SEM from 2-4 biological replicates.

### Divergent kinetics is independent of human IFNλ system

Our data suggests that in human cells, human IFNλ4 and its variants induce a distinct antiviral response compared with human IFNλ3. As previous work has demonstrated that IFNλ4 from different primate species have varying levels of antiviral activity (Bamford et al. 2018; Paquin et al. 2016), we next explored whether the distinct signalling kinetics that we observed were also species-specific. We first analysed the amino acid homology between IFNλ3, IFNλ4, IFNλR1 and IL10R2 in humans, chimpanzees and Rhesus macaques (**Fig 5A**). Results showed that although the various orthologues shared a high degree of homology (92-97%), there were differences that could affect activity given that even a single amino acid change can alter signalling as in IFNλ4 variants P70S and K154E. Given these genetic differences, we next tested the antiviral kinetics of non-human IFNλs. Firstly, we treated human HepaRG cells with human and non-human IFNλs as described in sFig2, by treating cells for 2h, 6h and 24h, and then removing the cytokines prior to infection with EMCV at 24h after initial stimulation **(Fig 5B)**. For these experiments we utilised non-human primate IFNλ3 or IFNλ4 proteins containing a C-terminal FLAG tag, which we characterised previously (Bamford et al. 2018). In these experiments we utilised IFNλ4 K154E as a model human IFNλ4 since it gave robust levels of detectable antiviral activity, with kinetics broadly similar to IFNλ4 WT. All IFNλs had antiviral activity against EMCV with chimpanzee IFNλ4 having greater activity than human and macaque IFNλ4 (**Fig 5C**), while human IFNλ3 had greater activity than macaque IFNλ3 (**Fig 5D**). Similar to human variants, the peak of IFNλ3 activity was delayed relative to all IFNλ4s. IFNλ3 Washing experiments demonstrated that like human IFNλ4, non-human primate IFNλ4 were more refractory to early removal than human or macaque IFNλ3 (**Fig 5C - D**). To determine if these characteristics also occurred in non-human cells, we repeated these experiments in the Rhesus macaque respiratory epithelial cell line LLCMK2 (**Fig 5E and F**). Results showed that all IFNλ4s had similar kinetics of antiviral activity but different levels of potencies as found in HepaRG cells. Washing following by immediate infection supported the initial washing experiments with IFNλ3 antiviral activity being more sensitive to early removal of cytokine (**Fig 5F and G**).

**Figure 5.**
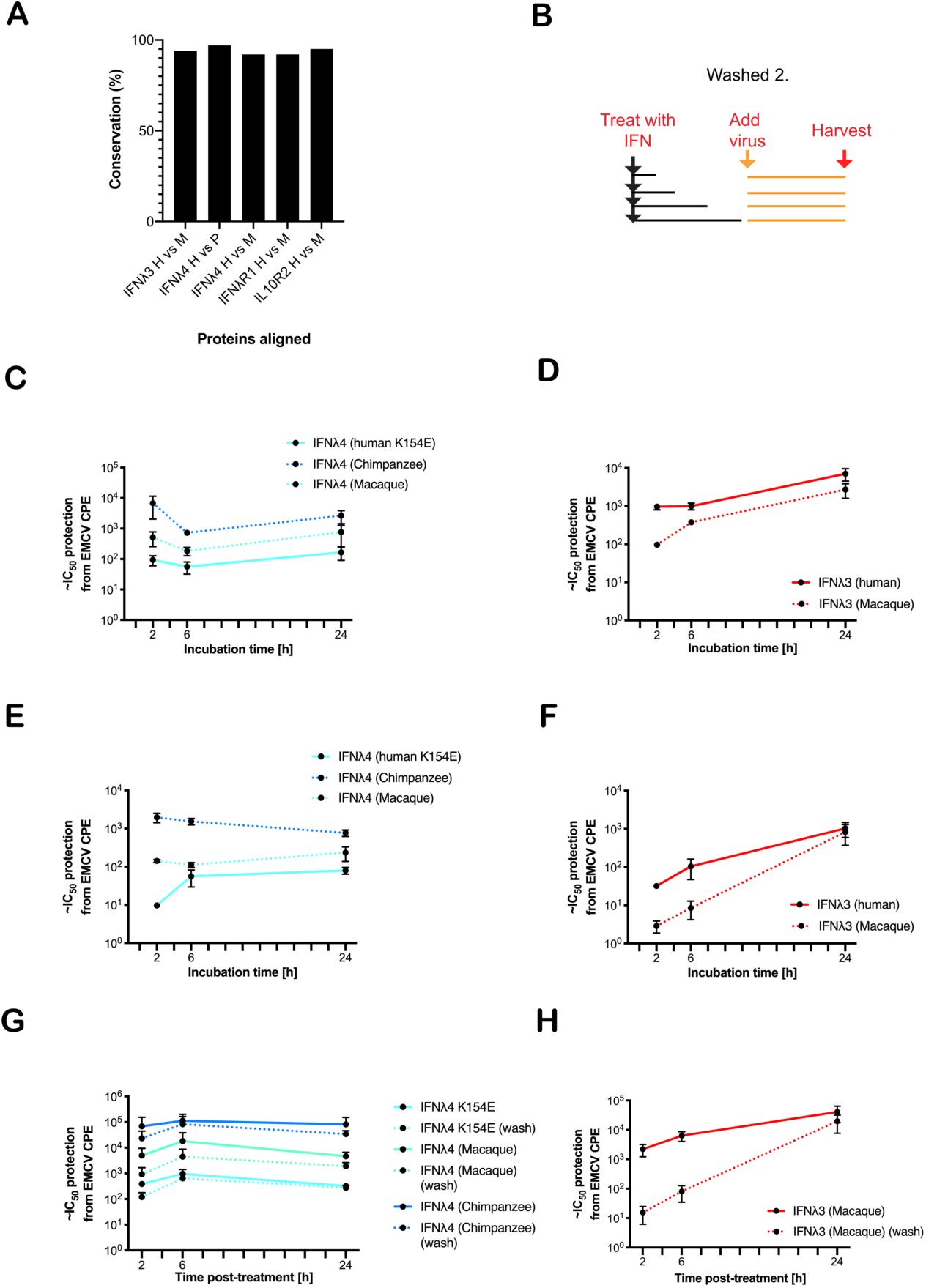
Kinetics of antiviral activity of non-human primate IFNλs. The percentage identity of IFNλ pathway proteins (IFNλ3, IFNλ4, IFNλR1 and IL10R2) between humans, chimpanzees and/or macaques was measured using BLAST (**A**). A washing/incubation protocol was used (**B**) and HepaRG (**C, D and G, H**) or Rhesus macaque LL-CMK2 (**E and F**) cells were pretreated with IFNλ4 (**C, E and G**) or IFNλ3 (**D, F and H**) for the indicated times prior to infection with EMCV. Times 24 (HepaRG) or 72 (LL-CMK2) hours post-infection antiviral activity was measured by CPE assay. Antiviral activity of IFNλs on HepaRG cells was measured using the alternative washing protocol (**G and H outlined in Fig 4**). Results are shown as mean ± SD from 4 biological replicates.

### IFNλ1 with receptor-interacting face mutations retain parental kinetics

Complex and dynamic interactions between cytokine ligands and their cognate receptors dictate the signalling output (Schreiber 2017). To probe further the molecular genetic basis of IFNλ kinetics we sought to mutate and disrupt the receptor binding faces of IFNλ hypothesising that these residues were most likely to be responsible for IFN kinetics. IFNλ4 is highly divergent when compared with IFNλ1-3 with ∼30% similarity detected suggesting that there are likely to be distinct molecular determinants of differential signalling contained within IFNλ4 compared to the other human IFNλs (Prokunina-Olsson et al. 2013; Hamming et al. 2013). To begin to identify those determinants, we constructed chimeric IFNλs between IFNλ4 and human IFNλ1. IFNλ1 was chosen as it is known to have similar kinetics to IFNλ3 (Marcello et al. 2006) but like IFNλ4, is N-linked glycosylated (Kotenko et al. 2003). Initially, comparison of differentially conserved amino acids in IFNλ4 (human and non-human primate) with IFNλ1-3 (human and macaque) identified a divergent receptor binding interface between these groups of IFNλs suggestive of distinct receptor interactions (**Fig 6A**). We focused on divergent, likely surface-exposed residues near relevant helices (A, D and F), and designed two chimeric IFNλs based on IFNλ1 containing candidate IFNλ4 residues from the IFNλR1-binding helix F (F), and the IL10R2-binding helices A and D (AD). An additional chimera with all three IFNλ4 binding helices was generated, termed ADF.

**Figure 6.**
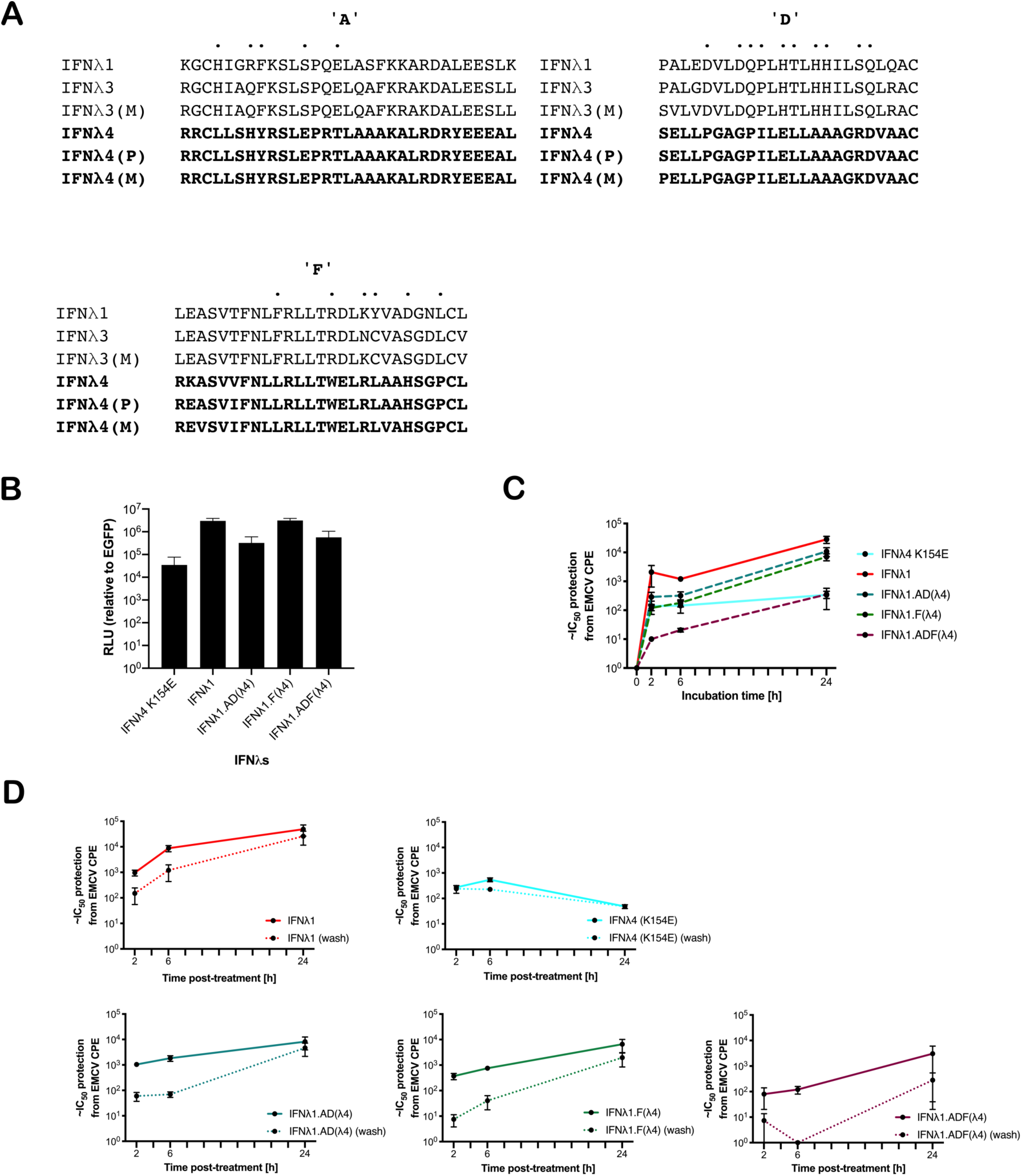
IFNλ1 receptor-interacting interface mutants retain their kinetics. IFNλ1/4 chimeras were generated based on critical differences in helices A, D and F identified by alignment and comparative approaches (**A**). Relative levels of IFNλs in supernatant by HiBiT assay following transfection of expression plasmids into HEK-293T cells measured at 48h after transfection (**B**). Effect of incubation time [2, 6 or 24h] (**C**) and washing [2, 6 or 24h, washed as hashed lines] (**D**) of antiviral activity in HepaRG cells against EMCV was calculated as outlined previously. Hatched line indicates limit of detection of that experiment. Error bars represent the mean ± SEM from 2-4 biological replicates.

We first confirmed that IFNλ1 and its chimeras were produced and released into the supernatant using a split-luciferase assay; this showed that chimeras incorporating helices A and D yielded reduced production (**Fig 6B**). To test their antiviral activity, HepaRG cells were pretreated for 2, 6 or 24 hours prior to EMCV infection with the WT IFNs and each of the indicated chimeras. Results showed that IFNλ1 had higher antiviral activity than IFNλ4 yet similar to IFNλ3. Additionally, IFNλ1 with IFNλ4 substitutions had reduced antiviral activity (IFNλ1>F>AD>ADF) (**Fig 6C and D**). To determine if the chimeras impacted IFN kinetics, HepaRG cells were pretreated for 2, 6 or 24 hours prior to EMCV with the WT IFNs and each of the indicated chimeras. The IFNs were either left for the duration of the infection or removed at the time of infection and infection was commenced either at time of cytokine removal or at 24h after initial incubation. Importantly, IFNλ1 kinetics were similar to IFNλ3 in HepaRG cells, with increasing activity over time and a delayed peak relative to IFNλ4 (**Fig 6C and D**). IFNλ1/4 chimeras had similar kinetic profiles as IFNλ1. Results revealed a that washing reduced the antiviral potency of all IFNs, IFNλ1 and all the chimeras were more greatly affected than IFNλ4. Taken with our antiviral activity results suggested that chimera F had reduced potency compared to IFNλ1, while the reduced activity of AD is likely due to reduced protein, and thus the impact on ADF is due to reduced amount and potency (**Fig 6B)**. However, despite alteration of the receptor interaction surfaces, the kinetics remain conserved similar to IFNλ1 (and IFNλ3), suggesting that these residues only modify the magnitude of the antiviral response and are not sufficient to alter the antiviral kinetics.

Altogether, our work described here demonstrates the distinct yet conserved antiviral kinetics of human and non-human primate IFNλ4 compared to other IFNλs.

## Discussion

Knowledge of the molecular signalling pathways stimulated by IFN binding is essential to understand immunity to infectious diseases, and could help develop more effective interventions. The dynamics of antiviral signalling is emerging as a physiologically-relevant and important topic and several groups have shown that type III IFNs have distinct slower but sustained signalling kinetics compared to type I IFNs (Marcello et al. 2006; Pervloraki et al. 2017; Pervolaraki et al. 2018). Very few studies have addressed whether different members of the type III IFN family also have a similar kinetics for the activation of STAT1, induction of downstream ISGs, and antiviral activity (Obajemu et al. 2017). Through several lines of genetic evidence, it appears that human IFNλ4 has non-redundant functions relevant to immunity compared to other IFNλs yet the determinants of this unique biology are poorly understood (Prokunina-Olsson et al. 2013; Terczyńska-Dyla et al. 2014). Additionally, there exist a number of naturally occurring functional variants of IFNλ4 that are known to impact potency (Terczyńska-Dyla et al. 2014; Bamford et al. 2018). In this work, we addressed whether IFNλ4 WT and its variants (e.g. P70S and K154E) have altered antiviral kinetics, in comparison to IFNλ1 and IFNλ3. By comparing IFNλ4 signalling and antiviral activity in two cell lines from two distinct organs, we were able to identify conserved and variable features of IFNλ4 and IFNλ3 signalling that demonstrated distinct antiviral kinetics, consistent with recent studies (Obajemu et al. 2017). Critically, we also show that common (P70S) and rare (K154E) human variants predominantly impact the magnitude of IFN signalling but not the kinetics of that response, and these dynamics are largely conserved in non-human primate IFNλs and their cognate cell lines.

Comparison of IFN activity across variants is notoriously challenging given the need for input normalisation and relevant processing. To circumvent these issues, we produced IFNλ in human cells (HEK-293T) and normalised for input IFNλ using a C-terminal “split luciferase” “HiBiT” tag system. Interestingly, using normalized amounts of protein released into the supernatant of transfected cells, we detected different potencies for each IFNλ, consistent with our previous work (Bamford et al. 2018). In general, IFNλ3 had greater antiviral potency than WT IFNλ4 in both human liver- and gut-derived cell lines. WT IFNλ3 induced stronger and more prolonged STAT1 phosphorylation, higher magnitude of ISG induction and a stronger antiviral effect than WT IFNλ4, which induced a lower and more transient response. The IFNλ4 K154E variant displayed potency that was more similar to IFNλ3 and shows that, at least for one rare variant, human IFNλ4 has the potential to have significant stimulatory effects. Considering the dynamics of the response, we show clear differences between IFNλ3 and IFNλ4 variants antiviral activity over time. These observations are consistent with previous work on IFNλ4 WT kinetics (Obajemu et al. 2017). Interestingly, IFNλ3 and IFNλ4 showed differential characteristics by limiting their contact time with target cells, suggestive of different interactions with receptor complexes. This observation requires more detailed biochemical and cell biology analysis, preferably using purified proteins and receptor molecules that would allow measurement of binding affinities, on-off rates, and their effects on receptor trafficking, for the IFNλ family. Such analysis was beyond the scope of our study primarily due to the inherent technical difficulties of preparing significant quantities of soluble, correctly-folded IFNλ4.

The interaction between IFNλs and their receptor complexes remains poorly understood, although several crystal structures of IFNλ proteins, with the exception of IFNλ4, in the presence and absence of its heterodimeric receptor complex IFNλR1 and IL10R2 have been solved (Gad et al. 2009; Mendoza et al. 2017). There is reason to believe that IFNλ4 is likely to interact differently with its receptors based on amino acid sequence alignments (Prokunina-Olsson et al. 2013; Hamming et al. 2013). IFNλ4 and IFNλ1/2/3 share only ∼30% homology, with highest levels found in the IFNλR1-binding ‘helix F’. Aside from helix F, IFNλ4 differs considerably compared to the other IFNλs, including other receptor binding helices, such as helix D that binds IL10R2. To test the contribution of IFNλ4 receptor interactors in and around helices A, D and F, we constructed chimeras using IFNλ1 as a reporter for antiviral activity into which we inserted predicted receptor binding domains from IFNλ4. These IFNλ1/IFNλ4 chimeric displayed similar kinetic profiles as IFNλ1 and IFNλ3 although differences in production and potency were noted. This suggests that the molecular determinants that regulate binding kinetics may not lie solely in the putative surface-exposed receptor-binding interfaces that we tested. IFNλ4 differs in structural capacity to IFNλ1/3, which may not be captured in our chimeras, and further differences are observed in other helices that may play roles in signalling. A possible explanation for these differences could be due to differing stabilities for each of the IFNλs. The stability of each IFNλ has not yet been tested but could provide insight into how each family member achieves its maximal activity. However, as most of our assays were performed in relatively short time frames (2-6h) it seems unlikely that IFN stability played a role in the differences we observed and is more likely that IFNλ4 interacts and activates the receptor more rapidly, likely through binding more strongly analogous to type I IFNs (Schrieber, 2017).

An important aspect of our work is that the differences we detected between IFNλ3 and IFNλ4 in antiviral kinetics were conserved in non-human species, through analysis of chimpanzee and macaque IFNλ4 and macaque IFNλ3 in human and macaque cell lines. This is important because compared to other primates, humans appear to have evolved unique IFNλ4 features relevant for outcome of infectious diseases like HCV (Prokunina-Olsson et al. 2013; Terczyńska-Dyla et al. 2014; Bamford et al. 2018). This finding would be consistent with the limited genetic differences between these species (>90% similarity). The fact that the kinetics are not unique to humans supports the hypothesis that alterations in IFNλ4 potency has been the dominant phenotype that our recent evolution has acted upon. It would be of interest to test further related IFNλ4, from distantly related mammals (Paquin et al. 2016).

Testing IFNλ kinetics in two cell lines allows us to assess conserved and divergent activities in hepatocytes and intestinal cells. IFNλs can signal in many tissues (Sommereyns et al. 2008), including the human gut (Perverolaki et al. 2017), and recent work has implicated variants in IFNλ4 in the outcome of enterovirus infection in the respiratory tract but which can infect the gut as well (Rugwizangoga et al. 2019). The role of IFNλ4 in intestinal cells up until now has been largely unexplored. While IFNλ3 and IFNλ4 can signal in both cell types, we show clear differences in potency of human IFNλ4 variants, consistent with our previous work in hepatocytes. Comparing the induction of IFNλ signalling in HepaRG and T84 cells suggested that the hepatocyte cell line was more sensitive to IFNλs yet to draw any conclusions, primary liver and intestinal cells or organoids from several individuals should be tested. Nevertheless, we observed consistent kinetics differences of IFNλ4 compared to IFNλ3 in both cell lines.

Our work has several implications, most importantly those relating to the conserved differences between IFNλ3 and IFNλ4. Compared to type I IFNs, IFNλs have been defined partially by their slower, sustained signalling kinetics (Marcello et al. 2006; Pervolaraki et al. 2018). IFNλ4 has several unique features, including its association with certain diseases, transcriptional suppression, and evolution in humans, which suggests a degree of specialisation. Unlike other IFNλs, IFNλ4 appears to signal more like type I IFNs despite utilising IFNλR1 and IL10R2. Thus, IFN kinetics may not solely lie in receptor biology but in the interactions between cytokine and receptor. We hypothesise that one outcome of the kinetics of IFNλ1-3 outlined here where activity is dependent on time and local concentration, would be a more tunable strategy, which may have “adaptive” potential for mucosal surfaces where more robust IFN activities may have pathogenic effects. Whether the unique kinetics of IFNλ4 would provide additional non-redundant therapeutic benefit over other IFNλs remains to be explored.

In conclusion, we provide further evidence of the functional divergence of IFNλ4 compared to other IFNλ proteins supporting the continued investigation into the causes and consequences of such distinctive signalling on the human immune system, which may be exploited for therapeutic gain.

## Acknowledgements

This work was funded by the UK Medical Research Council (https://mrc.ukri.org/) (MC_UU_12014/1) (JMcL). MLS and SB were supported by research grants from the Deutsche Forschungsgemeinschaft (DFG): (Project number 240245660 and 278001972 to SB and 416072091 to MLS. CG was supported by the China Scholarship Council and the Landesgraduiertenfoerderung fellowship from Heidelberg University. The funders had no role in study design, data collection and analysis, decision to publish, or preparation of the manuscript. Finally, we would like to thank the members of the McLauchlan, Boulant and Stanifer labs for helpful discussions; and Dr Lindsay Broadbent for critical reading of the manuscript prior to publication

---

**Supplementary Figure 1.**
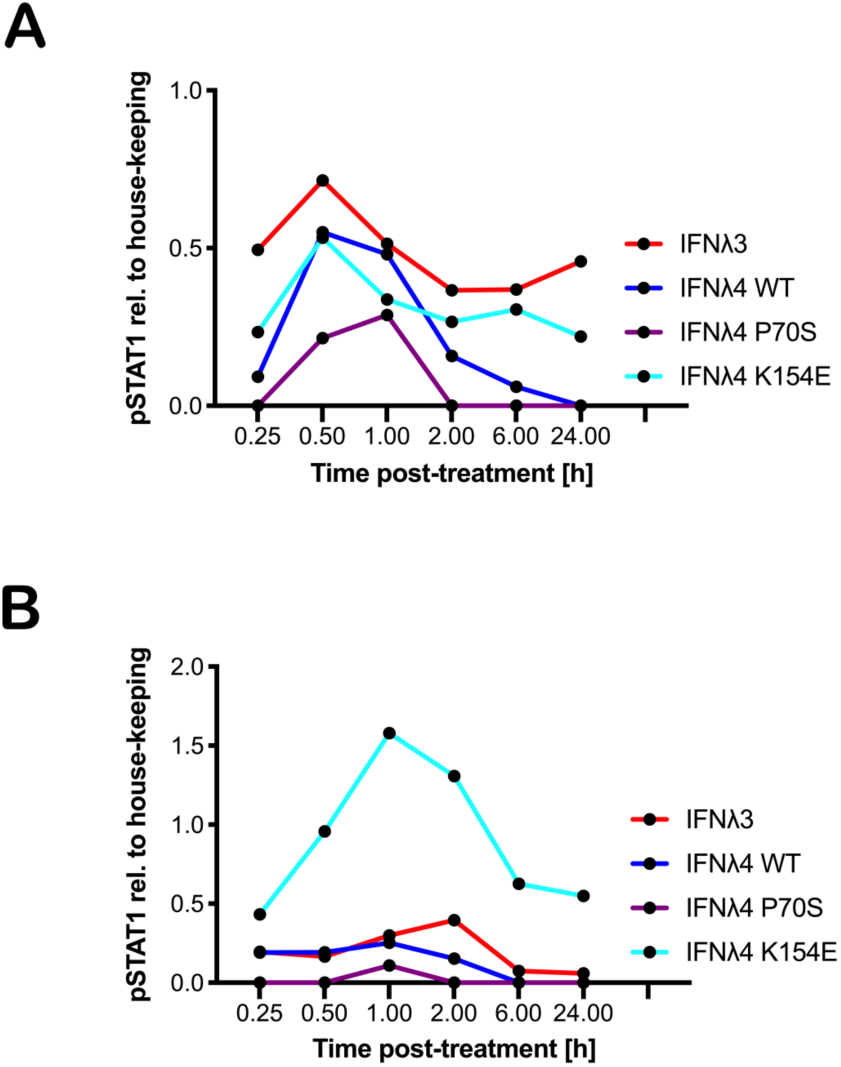
pSTAT1 quantification over time for IFNλs on liver and gut cells. Quantification of pSTAT1 from images in Fig 1 compared to house-keeping control and background levels was carried out by densitometry analysis for HepaRG (**A**) and T84 (**B**) cells (IFNλ3-HiBiT [red], IFNλ4-HiBiT: WT [blue], P70S [purple], and K154E [cyan].

**Supplementary Figure 2.**
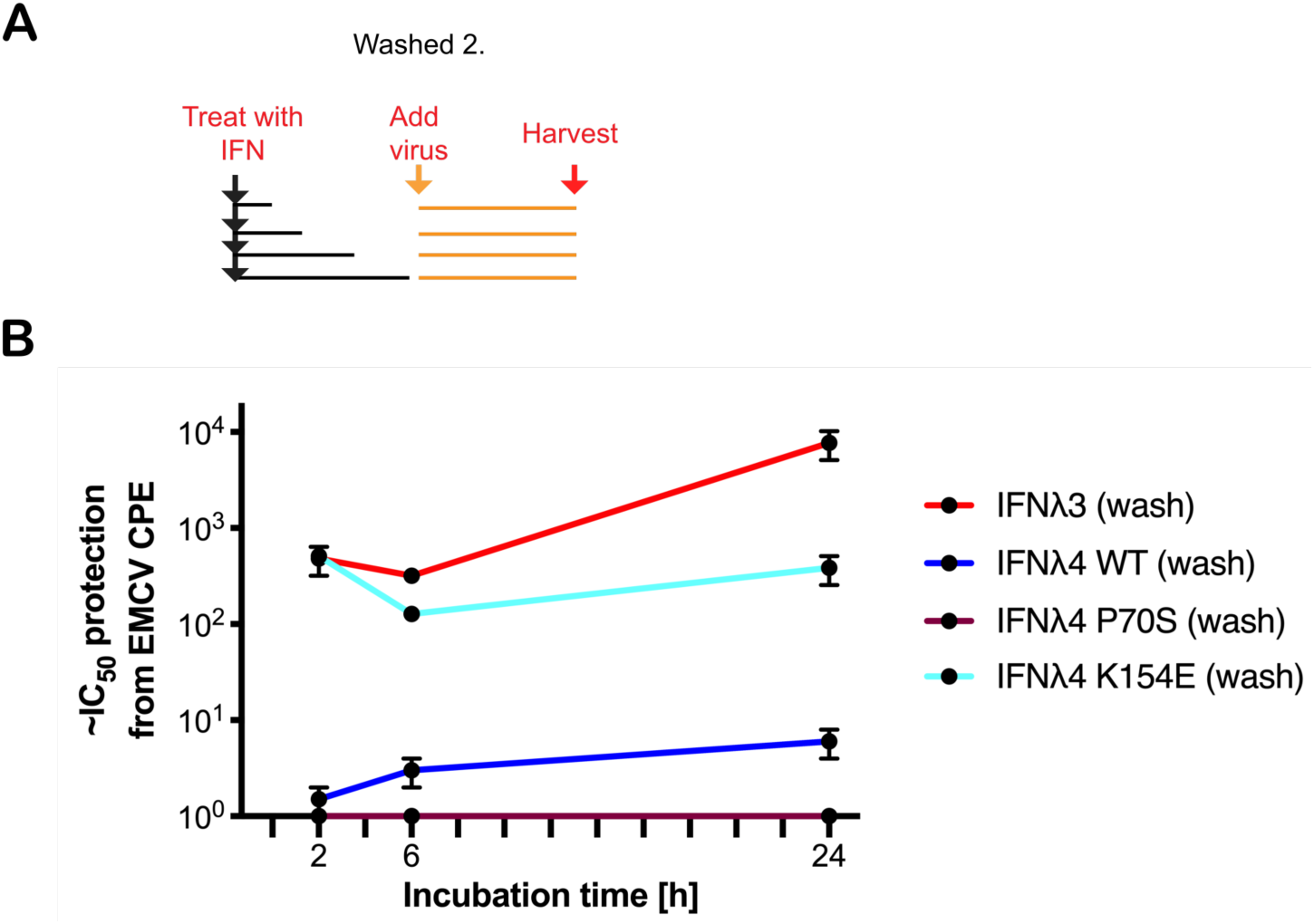
Effect of IFN incubation time on kinetics of human IFNλ variants. HepaRG cells were stimulated with IFNλs: IFNλ3-HiBiT (red), IFNλ4-HiBiT WT (blue), P70S (purple), and K154E (cyan) at indicated times (2, 6 or 24h) before supernatent was removed and rinsed with PBS before being replaced with fresh media not containing virus (**A**). Stimulated cells were incubated until 24h after IFNλ incubation prior to infection with EMCV and antiviral activity was read 24hpi (**B**). Error bars represent the mean ± SEM from 2 biological replicates.

## Materials and Methods

### Cell lines

HEK-293T (human embryonic kidney) and LLC-MK2 (Rhesus macaque respiratory epithelial cell line) were cultured in high glucose DMEM with 10% FCS and pen/strep. HepaRG.ISG15-EGFP (human hepatocyte-like cell line modified to express EGFP under the control of the endogenous ISG15 promoter (Bamford et al. 2018)) were cultured in complete William’s media with foetal calf serum (FCS) (10%), human insulin (4 µg/mL), hydrocortisone hemisuccinate (50 *µ*M) and pen/strep (1%) (HepaRG cells). T84 (ATCC CCL-248) colon carcinoma cells were cultured in a 50:50 mix of DMEM:F12 with 10% FCS and 1% pen/strep on collagen coated cell culture dishes. All cell lines were passaged routinely following PBS washing and trypsin-mediated detachment. Cell lines were routinely screened for Mycoplasma contamination and discarded if signs of contamination were detected.

### Viruses

Two viruses were used in this study: Ruckart strain of encephalomyocarditis virus (EMCV) and VSV. EMCV was produced in Vero cells following low MOI infection (MOI = 0.0001) and harvested between 1 and 2 days when extensive cytopathic effect was observed. EMCV infectivity was quantified by TCID_50_ and typically grew to titres of ∼10^8^/mL. VSV-luc was a kind gift from Sean Whelan (Washington University, St. Louis) and was produced and titrated as described in (DK et al. 2009; ML, DK, and SP 2011).

### Antibodies and reagents

Commercially available primary antibodies were mouse monoclonal antibodies recognizing β-Actin (Sigma #A5441), pSTAT1 (BD Transductions #612233), mouse anti-STAT1 antibody (3987, Abcam), or mouse anti-phospho-STAT1 antibody (29025, Abcam) and used at a 1:1000 dilution. Additionally, rabbit anti-Beta-Tubulin antibody (6046, Abcam) was also used (1:1000). For secondary antibodies: anti-mouse (GE Healthcare #NA934V), coupled with horseradish peroxidase (HRP) was used at a 1:5000 dilution (T84) or horseradish peroxidase-conjugated goat anti-rabbit IgG secondary antibody (A0545, Sigma-Aldrich) at 1:2000 dilution, or horseradish peroxidase-conjugated goat anti-mouse IgG secondary antibody (A4416, Sigma-Aldrich) at 1:2,000 (HepaRG).

### Molecular biology

Recombinant DNA technology was utilised to generate the IFNs for functional testing in this study, as previously described (Hamming et al. 2013; Bamford et al. 2018). The mammalian expression plasmids expressing HiBiT variants and human IFNλ3, as well as chimpanzee (*Pan troglodytes*) and Rhesus macaque (*Macaca mulatta*) IFNλ4 with a carboxy-terminal FLAG tag were described previously (Bamford et al. 2018). Rhesus macaque IFNλ3-FLAG was generated synthetically (GeneArt) with sequence corresponding to: XP_001086865.3 alongside WT IFNλ1-HiBiT or IFNλ1/λ4-HiBiT chimeras were constructed synthetically (GeneArt) with sequences from helices A, D and F as shown (**Fig 5**) and cloned into expression vector pC1 and sequenced confirmed by Sanger sequencing. Correct plasmids were purified by midiprep or maxiprep and quality and quantity determined by nanodrop prior to transfection. An EGFP expression plasmid prepared in identical conditions was used as a negative control throughout.

### IFN production

IFNλs were produced using the protocol described previously, which is capable of generating functional IFNλs(Bamford et al. 2018). Briefly, IFN expression plasmids were transfected into sub-confluent HEK-293T cell monolayers, which are hyporesponsive to IFNλ signalling due to very low expression of IFNλR1(Hamming et al. 2013). Lipofectamine 2000 was used to transfect IFNλ plasmids per manufacturer’s instructions. IFNλs were routinely generated in 6 well plates or 10 cm dishes, and 2 µg and 14 µg of plasmids were used, respectively. Lipofectamine 2000 (2 µl) was used per µg of plasmid. Plasmids were transfected into cells in Optimem for 16-18 hours, before changing media to growth media (10% FCS) until 2 days post transfection was reached. Conditioned media was harvested, clarified by centrifugation, aliquoted and immediately frozen at -80 in. Relative levels of IFNλs were estimated using the extracellular HiBiT split luciferase assay by virtue of their C-terminal HiBiT tag by incubating IFN preparations with assay reagents and measured by manufacturer’s instructions (Nano-Glo HiBiT Extracellular Detection system, Promega) using a luminometer.

### Interferon treatments

For qRT-PCR or immunoblotting experiments, IFN stimulation was achieved by incubating cell monolayers with IFNλ-containing conditioned media at a defined concentration to have equivalent HiBiT signal for each sample. Cells treated with IFNs were incubated for the indicated period of time before either being processed. A previous titration analysis indicated that a ∼1:2 - 1:4 dilution of IFNλ4-WT is enough to give a robust induction of ISGs for all variants (Bamford et al. 2018) and antiviral response while limiting the amount of conditioned media added to cells (<50% of total volume). Therefore, WT IFNλ4 was used at the standard and the levels of other IFNs were normalised to this by virtue of the HiBiT tag. Based on HiBiT assay measurements, the relative ratios of supernatent were: IFNλ4(WT):P70S:L79F:K154E:IFNλ3, ∼1:2:2:0.2:0.01. For the analysis of pSTAT1 levels by immunoblotting, 100,000 cells were seeded in 500 µl of growth media, into sterile rat-tail-collagen-coated (T84) or untreated (HepaRG) 24 well plates. To analyse ISG expression levels by qRT-PCR, 50,000 T84 cells were seeded in 500 µl DMEM/F12 into sterile rat-tail-collagen-coated 48 well plates or 1,000,000 HepaRG cells were seeded into 2 mL of growth media into 6 well plates. Cells were treated either with HEK293T cell supernatants containing either IF λ3 or different IFNλ4 variants (λ4 wildtype (WT),K154E, P70S) or GFP conditioned medium (Mock). Prior to treatment, media were removed, cells were rinsed once in PBS and then treated with each IFN diluted in their corresponding growth media to achieve an equal concentration (as determined by HiBiT) and added to the cells in 500-1000 *µ*l/well. Cells were then incubated at 37°C and 5% CO_2_ until harvest. Cells were harvested at 15 min, 30 min, 1 h, 2 h, 4 h, 24 h and 48 h post treatment for the analysis of pSTAT1 protein levels by immunoblotting, whereas total RNA was isolated from T84 cells at 2, 4, 8, 12 and 24 h post treatment for the analysis of ISG expression levels by qRT-PCR.

### Viral infections

For the EMCV anti-viral assays, 5,000 cells were seeded per well in a 96-well plate 24 hours prior to treatment. At the day of treatment IFNs were added in 2-fold serial dilutions to the cells 24 hours prior to infection. Following IFN treatment, EMCV (MOI = 0.3) was added to the cells and infection was scored by CPE 24hpi visually or by crystal violet staining. EMCV is highly cytopathic in certain cell lines and very sensitive to IFN. The reciprocal of the dilution giving ∼ 50% protection was used as a semi-quantitative measure of IFNλ conditioned media activity.

For VSV infection, T84 or HepaRG cells were seeded in a white bottom 96-well plate. Cells were pre-treated prior to infection as indicated time points and concentrations of IFN-λ3, IFN-λ4 and its variants K154E, P70S. VSV-luc (MOI =1) was added to the wells and the infection was allowed to proceed for 8 h. At the end and the infection, media was removed, cells were washed 1 × with PBS and lysed with Cell Lysis Buffer (Promega) at RT for 20min. The same volume of Steady Glo (Promega) was added to the cells and incubated for 15 min. Luminescence was read using Tecan Infinite M200 Pro.

### Immunoblotting

At the time of harvest, cells were rinsed once with PBS and then lysed with 1X RIPA buffer (150 mM sodium chloride, 1.0% Triton X-100, 0.5% sodium deoxycholate, 0.1% sodium dodecyl sulphate (SDS), 50 mM Tris at pH 8.0 supplemented with phosphatase and protease inhibitors (Sigma-Aldrich or Thermofisher) for 5-10 min at RT (T84) or ice (HepaRG). Cell lysates were collected and roughly equal amounts of protein were then separated by SDS-PAGE in a 10% (HepaRG) or 12% (T84) polyacrylamide gel, following boiling and reducing. Lysates were then blotted onto a nitrocellulose membrane (T84) or PVDF (HepaRG) by wet-blotting. Membranes were blocked with blocking buffer (5% BSA in TBS containing 0.1% Tween-20 (TBS-T) for 1 h at RT while shaking. Primary antibodies (1:1000 dilution) were diluted in blocking buffer and incubated overnight shaking at 4°C. The membranes were washed four times in TBS-T for 10 min at RT. Then, secondary antibodies were diluted in blocking buffer and incubated for 1 h shaking at RT. Membranes were again washed four times in TBS-T for 10 min at RT. HRP detection reagent (GE Healthcare) was mixed 1:1 and incubated at RT for 2-3 min or ECL substrate is added (Immobilon crescendo western HRP substrate, WBLUR0100, Merck). Membranes were then exposed to film and developed or visualised by chemiluminescence using the G:BOX Chemi gel doc Imaging System Instrument (Syngene). The detection of β-actin (T84) or β-tubulin (HepaRG) were used as loading controls. For quantitative analysis, pSTAT1 intensities of each immunoblot were quantified for each time-point using ImageJ or Image Studio Lite Version 5.2. For quantification with ImageJ, the background value (Mock) was manually subtracted from the calculated values. pSTAT1 levels were then determined relative to control.

### RT-qPCR

The total RNA was purified from lysed cells using the Nucleo Spin® RNA extraction kit (T84) by Marchery-Nagel (Catalog number 740955.50) according to the manufacturer’s instructions or (HepaRG) RNeasy Mini Kit (74106, Qiagen). RNA concentration was measured using the NanoDrop Lite spectrophotometer (Thermo Scientific). For T84 cells: 250 ng of total RNA was reverse transcribed into cDNA using the iScript™ cDNA Synthesis kit (BioRad Laboratories, Catalog number 1708891). The reaction contained a mixture of 1 μl Reverse Transcriptase, 4 μl Reaction Mix and 15 μl of RNA template in nuclease-free water. The newly synthesised cDNA was diluted 1:2 in RNase/DNase free water. The following qRT-PCR was performed using a Bio-Rad CFX96 Real-Time PCR Detection System. Per reaction 7.5 µl of SsoAdvanced Universal SYBR Green Supermix, 2 µl of 1:2 diluted cDNA, 1.7 µl of nuclease free water and 1.9 µl of either forward or reverse primers (2 µM) for the amplification of IFIT1 (fw: 5’
s-AAAAGCCCACATTTGAGGTG-3’; rev: 5’-GAAATTCCTGAAACCGACCA-3’), ISG15 (fw: 5’-CCTCTGAGCATCCTGGT-3’; rev: 5’-AGGCCGTACTCCCCCAG-3’), Viperin (fw: 5’-GAGAGCCATTTCTTCAAGACC-3’ and rev: 5’-CTATAATCCCTACACCACCTCC-3’) and Mx1 (fw: 5’-GGTCTATACCACACGCACAGA-3’; rev: 5’-ACTGGTTTCCTTTGCCTCGT-3’) were used. Data analysis was performed using the Bio-Rad CFX Manager 3.0. The expression of the targeted genes was then normalised to the housekeeping gene HPRT1 (fw: 5’-CCTGGCGTCGTGATTAGTGAT-3’; rev: 5’-AGACGTTCAGTCCTGTCCATAA-3’). For HepaRG cells: 1 μg of total RNA was reverse transcribed into cDNA using the High-Capacity cDNA Reverse Transcription Kit (Applied Biosystems, UK). The reaction contained a mixture of 1 μl Reverse Transcriptase, 9 μl Reaction Mix and 10 μl of RNA template in nuclease-free water. The newly synthesised cDNA was diluted 1:25 in RNase/DNase free water. The following qRT-PCR was performed using a RealTime Ready PCR Kit (Roche) and Taqman primer-primer-probe mixes. Each reaction mixture consisted of 10 μL of 2x LightCycler 480 Probes Master, 1 μL of 20x RealTime ready Assay with 4 μL PCR Grade H20 (total volume 15 μL). Template DNA, defrosted on ice, was first diluted 1:25 (v/v) with PCR grade H20 and then 5 μL diluted template added per reaction tube to the probes mastermix to give a final volume of 20 μL. Taqman assays (Cat no. 4331182) for: *IFIT1* (Assay ID: Hs03027069_s1), *ISG15* (Assay ID: Hs01921425_s1), *MX1 (*Assay ID: Hs00895608_m1), and *RSAD2/VIPERIN* (Assay ID: Hs00369813_m1) were used. The expression of the targeted genes was then normalised to the housekeeping gene *GAPDH* (Assay ID: Hs02786624_g1).

